# Effects of cLFchimera, a recombinant antimicrobial peptide, on intestinal morphology, microbiota, and gene expression of immune cells and tight junctions in broiler chickens challenged with *C. perfringens*

**DOI:** 10.1101/871467

**Authors:** Ali Daneshmand, Hassan Kermanshahi, Mohammad Hadi Sekhavati, Ali Javadmanesh, Monireh Ahmadian, Marzieh Alizadeh, Ahmmad Aldavoodi

## Abstract

The current study was conducted to investigate the effects of cLFchimera, a recombinant antimicrobial peptide (AMP), on various productive performance and gut health attributes of broilers experimentally challenged with *Clostridium perfringens* (Cp). Three hundred and sixty 1-day-old chickens were randomly allocated to 4 treatments of 6 replicates as follows: T1) unchallenged group fed with corn-soybean meal (CSM) without Cp challenge and additives; T2) challenge group fed with CSM and challenged with Cp without any additives; T3) peptide group challenged with NE supplemented with 20 mg cLF36/kg diet (AMP); T4) antibiotic group challenged with NE and supplemented with 45 mg antibiotic (bacitracin methylene disalicylate)/kg diet (antibiotic). Birds had free access to feed and water, sampling for villi morphology and ileal microbiota were performed on days 10 and 22, while jejunal section was sampled for gene expression of cytokines, tight junctions proteins, and mucin only on day 22. Results showed that AMP ameliorated NE-related lesion in the jejunum and ileum and reduced mortality in challenged birds compared to challenge group with Cp without any additives. Also, supplementing challenged birds with AMP improved growth performance and reconstructed villi morphology. While antibiotic non-selectively reduced the count of bacteria, AMP positively restored ileal microflora in favor of good bacteria (i.e. *Bifidobacteria spp.* and *Lactobacillus spp.*). AMP beneficially regulated the expression of cytokines, junctional proteins, and mucin in the jejunum of challenged birds with Cp. Since cLFchimera ameliorated NE lesion score, reduced mortality, improved productive performance and gut health attributes in chickens compared to challenged group and also were mostly similar with those of antibiotics and therefore, it could be concluded that this chimeric peptide can be a worthy candidate to substitute growth promoter antibiotics, while more research is required to unveil the exact mode of action of this synthetic peptide.

**Author summary:** Necrotic enteritis (NE) is a detrimental enteric disease in the poultry industry worldwide. The etiological factor of this disease is *Clostridium perfringens*, which is gram-positive anaerobic bacterium. This bacterium is common inhabitant of the intestine in lower counts (105), but it becomes pathogenic in higher counts and can secrete NetB toxin, which is the main cause of inducing NE in broilers. Due to the emergence of antibiotic-resistant bacteria, new generation of antimicrobial additives such as antimicrobial peptides (AMPs) has been introduced to the poultry industry. AMPs are small molecules with 12-50 amino acids having antibacterial activity. Recently, we extracted new AMP from camel milk, expressed in *E. coli*, refined and lyophilized to produce purified peptides. The current study investigated the effects of this peptide on prevention of NE in broilers. Results showed that AMP ameliorated lesion scores in the intestine and reduced mortality in challenged birds. AMP improved growth performance and reconstructed villi morphology in NE-challenged broilers. While antibiotic non-selectively reduced the count of bacteria, AMP positively restored ileal microflora. AMP beneficially regulated the expression of cytokines, junctional proteins, and mucin in the jejunum of NE-challenged birds.

## Introduction

Necrotic enteritis (NE) is well-known as a detrimental disease in the poultry industry, which results in production losses, increased mortality, reduced welfare of birds, and also increased risk of contamination of poultry products for human consumption [1]. The etiologic cause of NE is *Clostridium perfringens* (*C. perfringens*), a spore-forming Gram-positive bacterium, which is naturally inhabitant of farm animals gastrointestinal tract [2]. Antibiotics have been widely used to control NE in poultry farms, while the administration of growth-promoting antibiotics was extensively forbidden due to the rapid spread of antibiotic resistance as the main concern in human health [3]. The prohibition of using antibiotics in livestock industry has inspired researchers to search for safe substitutions for antibiotics and several additives have been introduced to the market, such as pro and prebiotics, essential oils, acidifiers, and antimicrobial peptides [4].

Antimicrobial peptides (AMPs) are endo-exogenous polypeptides comprised of less than 50 amino acids, characterized by cationic amphipathic properties, and produced by host defense systems or synthetically supplied to the diet in order to protect a host from pathogenic microbes [5, 6]. AMPs show broad-spectrum antimicrobial activities against various microorganisms, including Gram-positive and Gram-negative bacteria, fungi, and viruses [5]. These peptides are well-known for their roles as competent alternatives for antibiotics in farm animal production [7, 8, 9, 10, and 11]. The results of these studies demonstrated that AMPs could improve growth performance, nutrient digestibility and gut health, positively alter intestinal microbiota, and enhance immune function in pigs and broilers.

cLFchimera is a heterodimeric peptide designed to mimic two antimicrobial domains, Lactoferricin (LFcin) and Lactoferrampin (LFampin), which are present in the N1-domain of camel lactoferrin (cLF) [12]. More recently, the recombinant form of cLFchimera has been cloned and expressed in *E. coli* [12] and *L. lactic* [13] in our lab. The results of *in vitro* studies showed that cLF36 has antibacterial [12, 13 and 14] antiviral [15], and anticancer [16] properties. Furthermore, the results of an *in vivo* experiment showed that supplementing *E. coli* challenged broilers with cLFchimera improved villi morphology in the jejunum, restored microbial balance in the ileum, and improved gene expression of cytokines and tight junctions in the jejunum of challenged broiler chickens [17]. Therefore, the objective of the present study was to evaluate the effectiveness of cLFchimera as an alternative to growth enhancer antibiotics on performance and intestinal morphology, microflora, and gene expression of immune cells and junctional proteins in broiler chickens challenged with *C. perfringens*, as an animal model for infectious disease.

## Materials and Methods

### Ethics statement

All animal experiments conducted in the present study were in compliance with Iranian legislations in Agricultural Ministry, Deputy of Livestock and Veterinary Affairs (National Veterinary Organization, Iran). The ethic committee of Animal Care and Use of Ferdowsi University of Mashhad reviewed and approved the animal study protocols (Number 3/42449).

### Birds, treatments, and experimental design

A total of 360 1-day-old male chicks (Cobb 500) were purchased from a local hatchery, weighed and randomly assigned to 4 treatments with six replicates containing 15 birds in each replicate. Treatments were as follow: 1) unchallenged birds received a corn-soybean meal basal diet without AMPs, antibiotic, and Cp challenge; 2) challenge birds experimentally challenged with Cp; 3) birds experimentally challenged with Cp and supplemented with 20 mg peptide/kg diet (AMP); 4) birds experimentally challenged with Cp and supplemented with 45 mg antibiotic (bacitracin methylene disalicylate)/kg diet (antibiotic). All diets were in mash form and formulated to meet or exceed the minimum requirements of Cobb 500 (Table 1). Feed and water and were provided *ad libitum*. Chicks were reared in floor pens (1.1m × 1.3m) covered with wood shavings. Temperature and lighting programs were adjusted based on the guidelines of the Cobb 500 strain.

**Table 1.** Composition of experimental diets.

| Ingredient (%)<sup>1</sup> | Starter (0-10 days) | Grower (11-22 days) |
| --- | --- | --- |
| Corn | 56.81 | 58.16 |
| Soybean meal (44.0 %) | 36.01 | 34.80 |
| Soybean oil | 3.18 | 3.40 |
| Dicalcium phosphate | 1.79 | 1.65 |
| Limestone | 0.97 | 0.93 |
| Salt | 0.35 | 0.30 |
| Mineral-vitamin premix<sup>2</sup> | 0.50 | 0.50 |
| DL-Methionine | 0.17 | 0.15 |
| L-Lysine HCl | 0.22 | 0.12 |
| **Calculated nutrients** |  |  |
| AME (kcal/kg) | 3000.00 | 3080.00 |
| Crude protein (%) | 21.00 | 19.00 |
| Calcium (%) | 0.90 | 0.84 |
| Available phosphorus (%) | 0.45 | 0.42 |
| Sodium (%) | 0.16 | 0.16 |
| Methionine (%) | 0.50 | 0.47 |
| Methionine + cysteine (%) | 0.98 | 0.86 |
| Lysine (%) | 1.32 | 1.18 |

<sup>1</sup>Antibiotic (45 mg bacitracin methylene disalicylate/kg diet) and peptide (20 mg/kg diet) were added on top and thoroughly mixed.
<sup>2</sup> Added per kg of feed: vitamin A, 7,500 UI; vitamin D3 2100 UI; vitamin E, 280 UI; vitamin K3, 2 mg; thiamine, 2 mg; riboflavin, 6 mg; pyridoxine, 2.5 mg; cyanocobalamin, 0.012 mg, pantothenic acid, 15 mg; niacin, 35 mg; folic acid, 1 mg; biotin, 0.08 mg; iron, 40 mg; zinc, 80 mg; manganese, 80 mg; copper, 10 mg; iodine, 0.7 mg; selenium, 0.3 mg.

### AMP production

The AMP used in the present study was derived from camel lactoferrin (cLF) consisting of 42 amino acids, which were recently generated in our lab recently (for more details regarding the peptide cLF chimera production, please review previous papers [12, 13 and 14]. Briefly, preparation of recombinant plasmid vector was conducted through transforming recombinant expression vector harboring synthetic cLFchimera into DH5α bacterium [12, 13 and 14]. Next, the latter bacterial colonies were cultured to harvest plasmid extraction. Then, the recombinant vector was transferred into *E. coli* BL21 (DE3) as an expression host and cultured in 2 mL Luria-Bertani broth (LB) medium for overnight according to standard protocol [18]. In the next step, cultured materials were inoculated in 50 mL LB and incubated at 37°C with shaking at 200 rpm. Then, isopropyl-β-D-thiogalactopyranoside (IPTG) was added to a final concentration of 1 mM and incubated at 37°C for 6 h after IPTG induction. Periplasmic protein was collected at different times after IPTG induction (2, 4, and 6 h) according to the method described by de Souza Cândido et al. [19] and analyzed on 12% SDS-PAGE. To purify expressed peptide, Ni-NTA agarose column was used based on manufacturer’s instruction (Thermo, USA). The quality of purified recombinant peptide was assessed on a 12% SDS-PAGE gel electrophoresis, while the Bradford method was used to analyze the quantity of recombinant peptide. More recently, an *E. coli* expression system was developed in our laboratory that is able to produce 0.42 g/L of recombinant peptide. In the current study, 4 g peptide previously obtained from the recombinant *E. coli* were purified, lyophilized, and thoroughly mixed with 1 kg soybean meal and then supplemented to the relevant experimental diets.

### *C. perfringens* challenge

The method of Cp challenge was according to the method described in detail elsewhere [20], with some modifications. Briefly, on day 16, chicks in unchallenged groups were administered a single 1 mL oral dose of sterile phosphate-buffered saline (PBS) (uninfected) as a sham control, while challenged, peptide and antibiotic groups were orally challenged with 5,000 attenuated vaccine strain sporulated oocysts each of *E. maxima*, *E. acervulina*, and *E. tenella* (Livacox T, Biopharm Co., Prague, Czech Republic) in 1 mL of 1% (w/v) sterile saline. On days 20 and 21, birds in the challenge groups were orally inoculated with 1 mL *per os* the culture of *C. perfringens* (isolated from broilers meat [21], CIP (60.61) containing 10^7^cfu/mL in thioglycollate (Thermo-Fisher Scientific Oxoid Ltd, Basingstoke, UK) broth supplemented with peptone and starch. PCR analysis of inoculated *C. perfringens* in the current study was performed to confirm the presence of *netB* gene required for inducing NE in broilers according to Razmyar et al. [22]. The unchallenged groups received the same dose of sterilized broth.

### Growth performance

On days 10 and 22, body weight (BW) and feed refusal remaining feed of each pen were weighed to calculate the average daily gain (ADG), average daily feed intake (ADFI), and feed conversion ratio (FCR) over the specific and entire periods of experiment (0-10, 11-22, and 0-22 days of age). The feed conversion ratio for each period was readjusted based on the mortality data per pen per day, if any.

### Sample collection and lesion score

On days 10 and 22, 2 birds from each pen (12 birds/treatment) were randomly selected, euthanized by cervical dislocation, the viscera was excised, the intestine was discreetly separated from the whole viscera, and the adherent materials were precisely removed. The ileum was gently pressed to aseptically collect ileal content into sterile tubes for microbiological analysis. A section (about 5cm) from mid-jejunal tissues was meticulously separated for morphological analysis. A 2cm section from the mid-jejunum was detached, rinsed in cold phosphate-buffered saline (PBS), immediately immersed in RNAlater (Qiagen, Germantown, MD), and stored at −20°C for subsequent gene expression determination. On day 22, NE lesions of duodenum, jejunum, and ileum from 2 birds per pen were scored on a scale of 0 (none) to 6 (high) as described previously [23].

### Intestinal morphology

The method used to prepare samples for morphometry analysis was already explained by Daneshmand et al. [24]. Briefly, jejunal samples were stored in a 10% formaldehyde phosphate buffer for 48 h. Next, the samples were processed on a tissue processor (Excelsior™ AS, Thermo Fisher Scientific, Loughborough, UK), fixed in paraffin using an embedder (Thermo Fisher Histo Star Embedder, Loughborough, UK), and cut with a microtome (Leica HI1210, Leica Microsystems Ltd., Wetzlar, Germany) to a slice of 3cm. Then, the slices were placed on a slide and dehydrated on a hotplate (Leica ASP300S, Leica Microsystems Ltd., Wetzlar, Germany), and dyed the samples with hematoxylin and eosin. Finally, the dyed slices of jejunal were examined under a microscope (Olympus BX41, Olympus Corporation, Tokyo, Japan). A total of 8 slides were prepared from the jejunal segment per bird, and ten individual well-oriented villi were measured per slide (80 villi/bird). The average of slide measurements per sample was reported as a mean for each bird. Villus width (VW) was measured at the base of each villus; villus height (VH) from the top of the villus to the villus-crypt junction, crypt depth (CD) from the base of the adjacent villus to the sub-mucosa, the ratio of VH/CD and villus surface area were calculated.

### Microbial count

The methods used to count the populations of *E. coli*, *Clostridium spp.*, *Lactobacillus spp.*, and *Bifidobacterium spp.* in the ileal content were described elsewhere [25]. In summary, the ileal contents of a sample were thoroughly mixed, serially diluted 10-fold from 10^−1^ to 10^−7^ with sterile PBS, and homogenized for 3 minutes. Next, dilutions were plated on different agar mediums. Regarding the enumeration of bacteria, *Lactobacillus spp.* and *Clostridium spp.* dilutions were plated on MRS agar (Difco, Laboratories, Detroit, MI) and SPS agar (Sigma, Germany) and anaerobically cultured at 37°C for 48 h and 24 h, respectively. Black colonies of *Clostridium spp.* on SPS agar were counted. MacConkey agar (Difco Laboratories, Detroit, MI) and BSM agar (Sigma-Aldrich, Germany) were used to cultivate *E. coli* and *Bifidobacterium spp.* respectively, and incubated at 37°C for 24 h. All microbiological analyses were performed in triplicate, and average values were used for statistical analyses and results were expressed in colony-forming units (Log10 cfu/g of ileal content).

### RNA extraction and gene expression

The procedure of RNA extraction and gene expression was explained previously [26]. In summary, total RNA was extracted from chicken jejunum sampled on day 22 using the total RNA extraction kit (Pars Tous, Iran) following the manufacturer’s instructions. The purity and quality of extracted RNA were evaluated using an Epoch microplate spectrophotometer (BioTek, USA) based on 260/230 and 260/280 wavelength ratios, respectively. Genomic DNA was removed using DNase I (Thermo Fisher Scientific, Austin, TX, USA). The complementary DNA (cDNA) was synthesized from 1 µg of total RNA using the Easy cDNA synthesis kit (Pars Tous, Iran) following the manufacturer’s protocol.

Gene expression of two references (GAPDH and β-actin) and five targets (Interleukin-1 [IL-1], IL-6, mucin2 [MUC2], Claudin-1 [CLDN1], and Occludin [OCLN]) genes were determined by quantitative real-time PCR (qPCR) based on MIQE guidelines [27]. Each reaction was performed in a total volume of 20 μl in duplicate using an ABI 7300 system (Applied Biosystems, Foster City, CA) and 2× SYBR Green Real Time-PCR master mix (Pars Tous, Iran). Primer details are shown in Table 2. All primers were designed according to MIQE criteria [27] regarding amplification length and intron spanning. All efficiencies were between 90 and 110% and calculated R^2^ was 0.99 for all reactions. The method 2^−ΔΔCt Ct^ [28] was used to calculate relative gene expression in relation to the reference genes (GAPDH and β-actin).

**Table 2.**
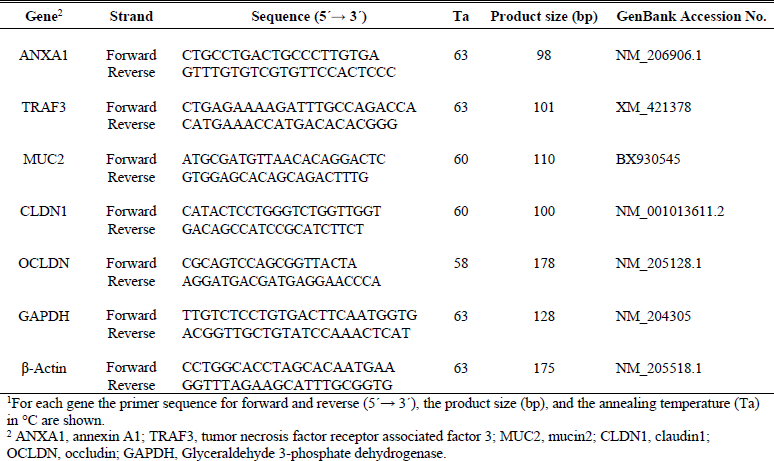
Sequences of primer pairs used for amplification of target and reference genes.^1^

### Statistical analysis

Data were statistically analyzed in a completely randomized design by ANOVA using the General Linear Model (GLM) procedure of SAS (SAS Inst., Inc., Cary, NC). Tukey’s test was used to compare differences among means of treatments, and P values < 0.05 were considered to be significant.

## Results

### Lesion score and mortality

Table 3 shows the effects of experimental treatments on NE-inducing lesion scores in different segments of the intestine and mortality rate of broiler chickens. Results showed that duodenum was mildly affected by NE, while additives had no significant effects on the recovery of this section from NE lesion. The results of lesion scores in the jejunum and ileum showed that the method of inducing NE was applied correctly. Peptide decreased (P < 0.05) lesions in the jejunum and ileum of birds compared to challenged group, while antibiotic intended (P > 0.05) to decrease lesions in the lower sections of the intestine. Birds fed antibiotic and peptide showed lower (P < 0.05) mortality rate compared to challenge group, while peptide group had no significant difference in comparison to unchallenged birds.

**Table 3.** Effects of treatments on necrotic enteritis lesion scores in broilers at 22 days of age.

| Treatments | Day 22 |  |  | Mortality<sup>2</sup> (%) |
| --- | --- | --- | --- | --- |
|  | Duodenum | Jejunum | Ileum |  |
| NC<sup>1</sup> | 0.000<sup>c</sup> | 0.000<sup>c</sup> | 0.000<sup>c</sup> | 0.000<sup>c</sup> |
| PC | 0.187<sup>a</sup> | 1.131<sup>a</sup> | 0.944<sup>a</sup> | 8.17<sup>a</sup> |
| AMP | 0.093<sup>b</sup> | 0.769<sup>b</sup> | 0.621<sup>b</sup> | 2.53<sup>bc</sup> |
| Antibiotic | 0.148<sup>ab</sup> | 0.955<sup>ab</sup> | 0.783<sup>ab</sup> | 3.44<sup>b</sup> |
| SEM<sup>3</sup> | 0.119 | 0.285 | 0.373 | 0.819 |
| P-value | 0.034 | 0.001 | 0.013 | 0.001 |

<sup>a-c</sup> Values within a column with different letters differ significantly ( $P < 0.05$ ).
<sup>1</sup> NC: negative control group received corn-soybean meal diet without challenge and additives; PC: positive control group received NC diet experimentally challenged with necrotic enteritis; AMP: PC received group supplemented with 20 mg antimicrobial peptide/ kg diet; Antibiotic: PC received group supplemented with 45 mg antibiotic (bacitracin methylene disalicylate)/ kg diet.
<sup>2</sup> Only mortalities shown necrotic enteritis symptoms.
<sup>3</sup> SEM: standard error of means (results are given as means ( $n = 12$ ) for each treatment).

### Broiler performance

Table 4 represents the effects of experimental diets on growth performance of broilers. Antibiotic fed birds showed higher (P < 0.05) ADG compared to challenged and unchallenged groups. Peptide also increased (P < 0.05) ADG when compared to the other group. However, its difference with unchallenged group was not significant. At d 22, additives decreased (P < 0.05) feed intake compared to challenged group, while their differences with unchallenged group were not significant. Interestingly, supplementing challenged chickens with peptide improved (P < 0.05) FCR compared to both challenged and unchallenged group at the end of the experiment (d 22), while birds received antibiotic showed better (P < 0.05) FCR compared to challenged group, but similar effect with those of unchallenged group.

**Table 4.** Effects of treatments on growth performance of broiler chickens from 0-22 days of age.

| Treatments | ADG<sup>2</sup> (g) |  |  | ADFI (g) |  |  | FCR (g/g) |  |  |
| --- | --- | --- | --- | --- | --- | --- | --- | --- | --- |
|  | 0-10 | 11-22 | 0-22 | 0-10 | 11-22 | 0-22 | 0-10 | 11-22 | 0-22 |
| NC<sup>1</sup> | 26.25<sup>b</sup> | 58.95<sup>b</sup> | 85.20<sup>b</sup> | 24.93<sup>a</sup> | 74.79<sup>b</sup> | 99.37<sup>b</sup> | 0.950<sup>a</sup> | 1.269<sup>b</sup> | 1.170<sup>b</sup> |
| PC | 26.15<sup>b</sup> | 55.91<sup>c</sup> | 82.06<sup>c</sup> | 23.77<sup>ab</sup> | 88.18<sup>a</sup> | 111.95<sup>a</sup> | 0.909<sup>a</sup> | 1.578<sup>a</sup> | 1.364<sup>a</sup> |
| AMP | 27.05<sup>ab</sup> | 60.29<sup>ab</sup> | 87.34<sup>ab</sup> | 22.61<sup>b</sup> | 72.23<sup>b</sup> | 94.85<sup>b</sup> | 0.836<sup>b</sup> | 1.199<sup>b</sup> | 1.086<sup>c</sup> |
| Antibiotic | 27.78<sup>a</sup> | 61.94<sup>a</sup> | 89.72<sup>a</sup> | 24.64<sup>a</sup> | 75.26<sup>b</sup> | 99.90<sup>b</sup> | 0.888<sup>ab</sup> | 1.215<sup>b</sup> | 1.113<sup>bc</sup> |
| SEM<sup>3</sup> | 0.318 | 0.446 | 0.644 | 0.422 | 1.401 | 1.577 | 0.0164 | 0.0257 | 0.0181 |
| P-value | 0.007 | 0.001 | 0.001 | 0.005 | 0.001 | 0.001 | 0.001 | 0.001 | 0.001 |

<sup>a-c</sup> Values within a column with different letters differ significantly (P< 0.05).
<sup>1</sup> NC: negative control group received corn-soybean meal diet without challenge and additives; PC: positive control group received NC diet experimentally challenged with necrotic enteritis; AMP: PC received group supplemented with 20 mg antimicrobial peptide/ kg diet; Antibiotic: PC received group supplemented with 45 mg antibiotic (bacitracin methylene disalicylate)/ kg diet.
<sup>2</sup>ADG: average daily gain; ADFI: average daily feed intake; FCR: feed conversion ratio.
<sup>3</sup>SEM: standard error of means (results are given as means of 6 pens of 15 birds/treatment).

### Jejunal villi morphology

The effects of treatments on jejunal morphology are shown in Table 5. On day 10, experimental diets had no significant effects on the morphometry of the intestine. Birds fed AMP and antibiotic had higher (P< 0.05) VH compared to non-challenged birds, while AMP had similar effect to that of non-challenged birds at 22 days of age. AMP enhanced (P < 0.05) villus surface area (VSA) compared to challenged and had similar effect in comparison to non-challenged group at 22 days of age, while antibiotic increased (P < 0.05) VSA compared to challenge group. Experimental diets had no significant effects on CD and VH/CD at 22 days of age.

**Table 5.** Effects of treatments on villi morphology (µm) in the jejunum of broiler chickens at 10 and 22 days of age.

| Treatment | Day 10 |  |  |  |  | Day 22 |  |  |  |  |
| --- | --- | --- | --- | --- | --- | --- | --- | --- | --- | --- |
| | VH<sup>2</sup> | VW | CD | VH/CD | VSA ($\mu\text{m}^2$) | VH | VW | CD | VH/CD | VSA ($\mu\text{m}^2$) |
| NC<sup>1</sup> | 621 | 188 | 144 | 3.29 | 367.7 | 1175<sup>a</sup> | 186<sup>a</sup> | 187 | 5.69 | 688.8<sup>a</sup> |
| PC | 592 | 192 | 125 | 3.09 | 356.6 | 827<sup>c</sup> | 153<sup>b</sup> | 201 | 5.04 | 396.9<sup>c</sup> |
| AMP | 681 | 194 | 138 | 3.52 | 414.2 | 1140<sup>ab</sup> | 187<sup>a</sup> | 171 | 6.06 | 671.5<sup>a</sup> |
| Antibiotic | 641 | 197 | 121 | 3.26 | 396.9 | 1017<sup>b</sup> | 174<sup>a</sup> | 180 | 6.50 | 557.0<sup>b</sup> |
| SEM<sup>3</sup> | 26.4 | 3.5 | 13.7 | 0.164 | 16.42 | 35.4 | 5.1 | 22.8 | 0.632 | 25.78 |
| P-value | 0.167 | 0.401 | 0.610 | 0.348 | 0.102 | 0.001 | 0.001 | 0.816 | 0.447 | 0.001 |

<sup>a-c</sup> Values within a column with different letters differ significantly ( $P < 0.05$ ).
<sup>1</sup> NC: negative control group received corn-soybean meal diet without challenge and additives; PC: positive control group received NC diet experimentally challenged with necrotic enteritis; AMP: PC received group supplemented with 20 mg antimicrobial peptide/kg diet; Antibiotic: PC received group supplemented with 45 mg antibiotic (bacitracin methylene disalicylate)/ kg diet.
<sup>2</sup>VH: villus height; VW: villus width; CD: crypt depth; VH/CD: the ratio of VH to CD; VSA: villus surface area.
<sup>3</sup>SEM: standard error of means (results are given as means ( $n = 12$ ) for each treatment).

### Bacterial colonization

Table 6 summarizes the effects of experimental diets on ileal bacterial populations. At d 10, antibiotic decreased (P < 0.05) the population of all bacteria compared to challenge and non-challenged groups, while AMP had similar effects to those of other treatments. Birds supplemented with antibiotic had the lowest (P < 0.05) population of all cultured ileal bacteria compared to both challenged and non-challenged groups at 22 days of age. Interestingly, AMP increased (P < 0.05) the population of *Lactobacillus spp.* and *Bifidobacterium spp.* and decreased (P < 0.05) the colonization of *E. coli* and *Clostridium spp.* in the ileum of chickens compared to challenge birds, while AMP had similar effects compared to non-challenged group at 22 days of age.

**Table 6.**
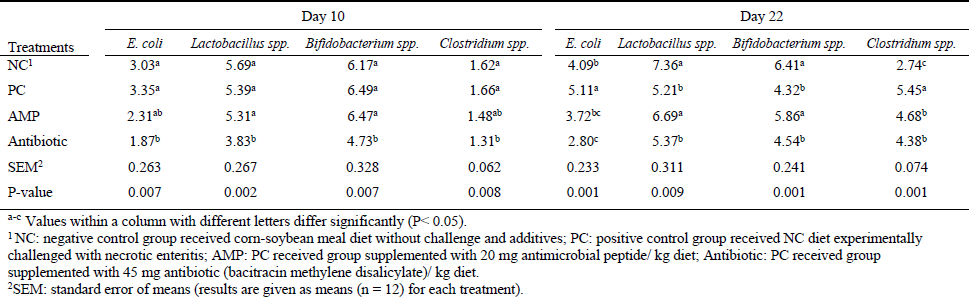
Effects of treatments on ileal microflora (log_10_ CFU g^−1^) in broilers at 10 and 22 days of age.

### Gene expression of immune cells and tight junction proteins

The effects of treatments on the expression level of immune cells and tight junction proteins are presented in Figure 1. While *C. perfringens* challenge increased (P < 0.05) TRAF3 and ANXA1 expressions, adding antibiotic and AMP to the diet reduced (P < 0.05) expression of these immune cells compared to challenged group and had similar effects to those of non-challenged birds.

**Figure 1.**
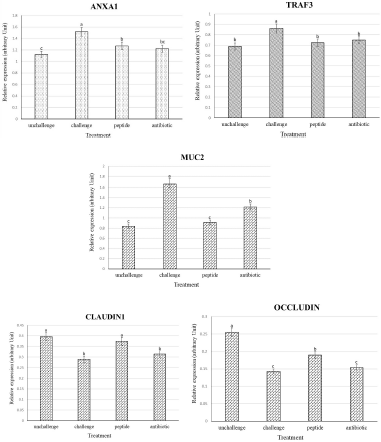
Effects of treatments on the expression of different genes in the jejunum of broiler chickens on day 22. Samples were analyzed using qPCR, and GAPDH and β-actin were used as the reference genes. Abbreviations as follows: ANXA1, annexin A1; TRAF3, tumor necrosis factor receptor associated factor 3; MUC2, mucin2; unchallenge, control birds received a corn-soybean meal basal diet without AMPs, antibiotic and necrotic enteritis (NE) challenge; challenge, control birds experimentally challenged with NE; peptide, birds experimentally challenged with NE and supplemented with 20 mg peptide/kg diet; Antibiotic, birds experimentally challenged with NE and supplemented with 45 mg antibiotic (bacitracin methylene disalicylate)/kg diet; The letters on the bar mean show significant difference (P < 0.05).

Antibiotic and AMP increased (P < 0.05) the expression level of MUC2 compared to the challenged group. While antibiotic had higher (P < 0.05) expression of MUC2 compared non-challenge birds, AMP had similar expression levels when compared to non-challenged groups. Antibiotic did not significantly affect gene expression of jejunal junctional proteins compared to non-challenged group and had similar effects to those of challenged birds. AMP improved (P < 0.05) expression patterns of CLDN1 and OCLN in the jejunum compared to challenged group.

## Discussion

Necrotic enteritis is still a global concern with drastic losses in poultry farms, mainly due to retarded growth performance, increased mortality, and veterinary costs [23]. The outbreak of disease and consequently economic losses have been more prominent in post-antibiotic era [23]. While the use of antibiotics has been banned in many parts of the world, such European Union, due to health concerns related to emerging antibiotic-resistance pathogens and also drug residues in poultry products, researchers have investigated for alternative additives to restrain NE. Recently, more attentions have inclined to AMPs due to their beneficial roles on health attributes and to prophylactic effects against pathogenic invasion [12, 13 and 17]. Therefore, the principal objective of the current study was to investigate the effects of antimicrobial peptide, cLFchimera, on various productive and health parameters in chickens experimentally challenged with NE.

Results of the current study showed that AMP decreased gut lesion and mortality induced by NE and also improved growth attributes in challenged chickens similar to antibiotic fed birds, which is in agreement with previous results [9, 10]. While most of the previous researches have studied the effects of AMPs in chickens in normal conditions, Hu et al. [29] demonstrated that supplementing broilers diet with AMP improved their weight gain and FCR under heat stress, which is in agreement with the current results. In another challenge study, Wu et al. [30] challenged weanling pigs with *E. coli* and supplemented the diet with AMP. They reported that AMP reduced the incidence of diarrhea post-challenge and improved weight gain and FCR compared to challenged group, which is similar to the present findings regarding the reduction in gut lesion and improvement in performance. Previous studies attributed the beneficial effects of AMPs on growth performance of chickens to their fundamental roles in maintaining microbial balance in the gut and consequently improvement in the intestinal morphometry [9, 10].

It has been well-documented that the villi play the critical roles in absorbing nutrients from the intestinal tract, which subsequently the morphometry of these villi can drastically affect the host’s performance and health [31]. In the present study, AMP significantly improved morphometry of villi in the jejunum of challenged chickens, similar to that of unchallenged group, which is in agreement with previous studies [9, 10]. It has been reported that AMPs extracted from pig intestine [32] and rabbit sacculus rotundus [33] enhanced jejunal villi characteristics in broiler chickens, which is consistent with the present results. Generally, in healthy conditions, provision of essential nutrients and microbial balance are two crucial factors affecting villi morphology [34]. On the other hands, in an infectious disease like NE, the most critical strategy in maintaining villi structure is the removal or leastwise elimination of the pathogens through providing antimicrobial additives and manipulating the intestinal microbiome [35]. Previous studies showed that antibiotic and AMPs could improve villi morphology and nutrient absorption and consequently increase growth performance in chickens under healthy and/or disease conditions by manipulating the intestinal microflora [9, 10 and 17].

The intestinal commensal microbiome interacts with the host through different processes, including nutrients absorption, villi morphology, intestinal pH, and mucosal immunity [36, 37]. In the current study, antibiotic reduced the colonization of all bacteria, while AMP significantly enhanced the beneficial bacterial populations (i.e. *Lactobacillus spp.* and *Bifidobacterium spp.*) and decreased the proliferation of opportunistic pathogen populations (i.e. *E. coli* and *C. perfringens*) in the ileum. In agreement with the present study, Tang et al. [7] and Ohh et al. [38] reported that AMPs significantly enhanced the population of beneficial bacteria and decreased the colonization of harmful ones in the ileum of piglets and broilers, respectively. The antimicrobial action of Bacitracin Methylene Disalicylate (BMD) involves blocking the bacterial ribosome subunits and subsequently impeding protein synthesis, which finally reduces the colonization of microbial community in the intestine [39]. Unfortunately, this antibiotic does not differentiate between beneficial vs. pathogenic bacteria and may perturb microbial balance in the intestine and deprive the host of benefits of microbes’ roles and products [40, 39]. There is no consensus on the mechanism by which AMPs influence bacterial colonization in the intestine, while two direct and indirect mechanisms have been proposed based on the physiological properties of peptides. The direct antimicrobial effect of AMPs has been attributed to different surface charges of peptides and pathogens [41]. In other words, AMPs possess positive charge contributing to electrostatically adhere to negatively charged bacterial membranes [42, 41]. This attachment can either destroy the bacterial membranes through physical disruption or penetrate the bacterial cytoplasm without exerting any damage to the lipid layer [43, 41]. Imported AMPs may interfere with intracellular signaling pathways like nucleic acids synthesis, enzyme activity, and protein biosynthesis [42, 43]. In the indirect mode, AMPs might manipulate the microbial community of the intestine in favor of the colonization of beneficial bacteria (e.g. *Lactobacillus spp.* and *Bifidobacterium spp.*) and beneficially affect the host health through various physiological mechanisms (e.g. competitive exclusion, secretion of short-chain fatty acids, activation of intestinal immune system, etc.) [42]. Previous findings suggested that cLF36 could attach to the bacterial membrane through electrostatic interactions and physically disrupt bacterial bilayer membranes [12, 13 and 14]. In line with the previous reports [44], the current results demonstrate that AMP can selectively prevent the bacterial growth in the intestine of *C. perfringens* challenged chickens, which may prove the competitive advantage of cLchimera compared to antibiotics. Furthermore, previous research reported that the antimicrobial activities of AMPs against pathogens in the intestine might alert host immune system to fight against invading agents [45, 46].

Mucosal immunity plays an important role in host defense against pathogens [47]. At the intestinal level, epithelial cells express pattern recognition receptors (PRRs), which can recognize molecular origins found on most classes of microbes called microbe-associated molecular patterns (MAMP). The recognition of MAMP by PRRs would lead to the stimulation of immune systems [48, 49]. Toll-like receptors (TLRs) are considered the most important PRRs which can recognize MAMP and facilitate the initiation of immune response against pathogen invasion [50, 51]. Furthermore, host immune system hires both pro-and anti-inflammatory cytokines to fight against invading pathogens and restore mucosal homeostasis [52, 53, and 54].

Pro-inflammatory cytokines like IL-1β, TNF-α, and IL-12 stimulate the body’s defense through immune cell differentiation, proliferation, apoptosis, and NO production [55]. In broiler chickens, when TLR4 engages to MAMP, it transmits the information to the cytoplasm of the phagocytes, which in turn leads to expression of cytokines [56, 57]. The controversial results have been reported by different research groups regarding the effects of *C. perfringens* challenge on gene expression of TLR4 in the intestine of broiler chickens. For instance, while some researchers reported that *C. perfringens* upregulated the TLR4 gene expression in the intestine of chickens [58, 59], other investigators reported no apparent alteration of the TLR4 gene expression in *C. perfringens* challenged chickens [60]. Therefore, in the current study, we decided to analyze the gene expression of TRAF3, which is one step ahead of TLR4 activation in order to overcome the possible interference of other immune cells [61]. TRAF3 is a cytoplasmic protein that controls signal transduction from different receptor families, especially TLRs [61]. Following the activation of TLR4 with pathogen attachment, TRAF3 is recruited into signaling complexes, and its activation increases vital pro-inflammatory cytokines production [62, 63]. Results of the present study showed that while *C. perfringens* challenged upregulated the expression of TRAF3 in the jejunum of chickens, antibiotic and AMP significantly decreased the expression of this cytokine in the challenged birds. To the best of our knowledge, this is the first experiment that reports the expression of TRAF3 under AMP and antibiotic treatments in *C. perfringens* challenged chickens, while there is consistency between the current results and previous ones regarding the effects of *C. perfringens* challenge on TRAF3 expression [62, 64].

On the other hand, excessive and long-term production of pro-inflammatory cytokines might result in the gut damage and high energy demand [55]. To prevent the adverse effects of extra pro-inflammatory cells, pro-resolving mediators such as ANXA1 are released into the epithelial environment to orchestrate clearance of inflammation and restoration of mucosal homeostasis [65, 52). ANXA1 is a 37 kDa calcium-and phospholipid-binding protein expressed in the apical and lateral plasma membrane in the intestinal enterocytes that facilitates resolution of inflammation and repair [66]. ANXA1 applies several mechanisms to induce anti-inflammatory effects. Primary mechanism include suppressing the release of pro-inflammatory cytokines like IL-1β and TNF-α in the intestinal mucosa, inhibiting leukocyte migration and monocyte adhesion to vascular endothelium, impairing neutrophil, and eventually promoting apoptosis of inflammatory cells [67, 68, 69]. In the current study, *C. perfringens* upregulated the expression of ANXA1 in the jejunum of challenged chickens, which is in agreement with previous report [62], while antibiotic and AMP significantly decreased the gene expression of this cytokine, which is firstly reported herein. While there is no well-documented evidence to explain the results of cytokines expression, it could be inferred that antimicrobial activities of antibiotic and AMP in the current study resulted in the reduction of invading pathogens (based on abovementioned microbial results) in the intestine of challenged birds and possibly downregulating the expression of cytokine-producing immune cells and finally a decrease in pro-and anti-inflammatory cytokines. While the same mechanisms have been proposed separately for the effects of BMD [70] and other AMPs [11] on cytokines expression in *C. perfringens* challenged chickens, no comprehensive mechanism has been reported by the time of preparing this paper. Along with the crucial roles in immune system, it has been shown that cytokines can affect junctional proteins and intestinal leakage [71].

The epithelial barrier consists of tight junction proteins forming the primary lines of defence against wide range of stimuli from feed allergens to commensal and pathogenic bacteria [72, 73]. The disruption of these proteins may result in increasing the intestinal permeability to luminal pathogens [72, 73]. Previous studies showed that *C. perfringens* might attach to the junctional proteins in order to form gaps between the epithelial cells and disrupt the intestinal integrity [73; 74]. In the present study, NE challenge reduced the jejunal gene expression of OCLN and CLDN1, which is in agreement with previous studies [74, 75], while AMP significantly upregulated the expression of these genes in the challenged birds, and antibiotic had no significant effect on the gene expression of junctional proteins. In agreement with the current findings, previous reports demonstrated that AMPs could increase the expression of junctional proteins in different challenge conditions [76, 17]. Previous studies showed that tight junction proteins, especially CLDN1 and OCLN, have a specific region (i.e. ECS2) containing a toxin-binding motif, NP (V/L)(V/L)(P/A), that is responsible for binding to *C. perfringens* [77, 73]. Following attachment to junctional proteins, *C. perfringens* could digest these proteins [78] and open the intracellular connection between adjacent epithelial cells resulting in more penetration of pathogens to deeper layers of lamina propria and transmitting to other organs [79 73]. While no exact mechanism has been recognized for the inhibitory effects of AMPs on *C. perfringens* regarding junctional proteins, two acceptable theories have been suggested. In the first theory, it has been suggested that AMPs could directly switch on the expression of regulatory proteins (i.e. Rho family) in the intestine of challenged mice that consequently upregulated the expression of tight junction proteins and ameliorated leaky gut [80, 76]. The second theory attributed the beneficial effects of AMPs on tight junctions to their indirect roles in manipulating microflora populations in the intestine. In detail, previous studies showed that the intestinal commensal bacteria like Bifidobacteria and Lactobacilli secrete butyric acid that regulates epithelial O_2_ consumption and stabilization of hypoxia-inducible factor, a transcription factor protecting the epithelial barrier against pathogens, resulting in higher expression of junctional proteins [81, 82]. As previously discussed in the current study, cLFchimera decreased the number of *C. perfringens* and increased the population of Bifidobacteria and Lactobacilli in the intestine. Therefore, it can be hypothesized that AMP in the current study upregulated the expression of junctional proteins through both reducing the number of *C. perfringens* and inhibiting proteins disruption by bacterial toxins, and increasing the concentration of butyric acid by increasing the count of butyrate-producing bacteria like Bifidobacteria and Lactobacilli. Surprisingly, antibiotic did not change the expression of CLDN1 and OCLN in the jejunum of challenged chickens, while it could be expected that antibiotic upregulated the junctional proteins due to the antibacterial nature of antibiotics. In line with the current results, Yi et al. [76] reported that antibiotics might not affect the gene expression of junctional proteins of the epithelial cells after pathogen removal, maybe because of controlling the microbial balance in the intestine.

Along with junctional proteins, the luminal mucus layer comprising of mucins plays a defensive role against invasive pathogens [83]. MUC2 widely expresses in goblet cells and secretes into the intestinal lumen to stabilize mucosal layer [83, 84]. Any damage to the mucosal layer and/or the interaction between MAMPs and PRRs stimulates the expression of MUC2 to secrete more mucin and prevent further destruction [84, 85]. In the current study, *C. perfringens* challenged significantly increased the expression of MUC2 in the jejunum, which is in agreement with the results of previous studies [86, 87]. On the other hand, antibiotic and AMP significantly downregulated the expression of this gene, while the results for AMP was similar to those of unchallenged group. According to the bacterial results in the present study, it could be inferred that the inhibitory effects of AMP on the population of *C. perfringens* and *E. coli* might reduce the colonization of these bacteria in the intestine, decrease the destruction of mucosal layer, and subsequently lessen the expression of MUC2, while the exact mechanism has not been revealed yet.

In conclusion, results of the current study propose that cLFchimera, an antimicrobial peptide originated from camel milk, could reduce mortality and attenuate NE-induced lesions resulted in better growth performance, recovery of villi morphology in the jejunum, and restoration of the ileal microflora in NE-imposed chickens. Furthermore, supplementing *C. perfringens* challenged birds with cLFchimera beneficially regulated the gene expression of cytokines to boost the immune system against co-inocculation of *Eimeria spp*. and *C. perfringens* in NE challenge model. Eventually, cLFchimera significantly repaired the intestinal mucosal layer and barrier functions in *C. perfringens* challenged chickens through positively manipulating the expression of MUC2 and tight junctional proteins. Therefore, according to the desired results obtained in the present study, cLFchimera can be nominated as a candidate for replacing growth promoter antibiotics against NE in chickens, while further studies may find other favourable effects of this AMP.

## Acknowledgments

The authors would like to thank Ferdowsi University of Mashhad, Iran, for their technical support and facility provision of this study (Number 3/42449).

## Author Contributions

### Conceptualization

Hassan Kermanshahi, Mohammad Hadi Sekhavati, Ali Javadmanesh.

### Formal analysis

Ali Daneshmand, Ali Javadmanesh

### Investigation

Ali Daneshmand, Mohammad Hadi Sekhavati, Ali Javadmanesh.

### Methodology

Ali Daneshmand, Monireh Ahmadian, Marzieh Alizadeh, Ahmmad Aldavoodi.

### Project administration

Hassan Kermanshahi, Mohammad Hadi Sekhavati.

### Supervision

Hassan Kermanshahi, Mohammad Hadi Sekhavati.

### Writing – original draft

Ali Daneshmand, Mohammad Hadi Sekhavati.

### Writing – review & editing

Ali Daneshmand, Hassan Kermanshahi, Mohammad Hadi Sekhavati

